# Establishing genetically controlled, closed colonies of an ascidian

**DOI:** 10.64898/2026.01.29.702678

**Authors:** The Ciona bio-resource consortium

**Author notes:** Aitsu Marine Station, Kumamoto University, 6061 Aitsu, Matsushima, Kami-Amakusa City, Kumamoto, 861-6102, Japan.

## Abstract

Recent technological advances have made many “non-model” organisms accessible for experimental studies. However, reference inbred strains were not necessarily available, especially in marine invertebrates, and genetic background of organisms used for experiments are often non-uniform. This situation potentially affects experimental reproducibility. Although ascidians, *Ciona intestinalis* (type A, or *C. robusta*), are a widely used marine animal for many areas of experimental biology including developmental studies, no reference strains have been obtained despite extensive efforts. As an alternative way to improve reproducibility, we have established and maintained ascidian colonies through intra-population breeding every year from 2016 to 2020, and monitored genomic variants of these colonies. This method does not reduce genetic variations but instead manages and monitors genetic variations in the colonies, providing an easy and cost-effective way of increasing experimental reproducibility. Furthermore, we recently upgraded these genetically isolated, closed colonies that were re-established every year, and have maintained them for more than three years only through intra-population breeding and occasional back-cross using cryopreserved sperm. Genetic variants that we revealed using 3.7 tera-bases of sequence data will help to design future experiments in this species. Our data also show that two wild-populations, which were used to establish the colonies, have maintained distinct genetic backgrounds, although their habitats are directly linked to the Pacific Ocean and only 170 km apart. More importantly, genetic information regarding these colonies will undoubtedly improve experimental reproducibility and traceability, and our method will provide a realistic solution for performing reproducible experiments using non-model organisms.

## Introduction

Recent technological advances have made many animal species accessible, which formerly were rarely used for experiments. For example, next generation sequencing technology has made it easier to decode entire genome sequences, and CRISPR-CAS9 technology has made it possible to test gene functions (Wang and Doudna 2023). However, for animals that have only recently begun to be used for experiments, especially marine invertebrates, no inbred reference strains are available. This situation potentially affects reproducibility of experiments, especially in animals with high heterozygosity. Indeed, the influence of genetic background on phenotype is well-documented in model organisms. Notable examples include *Apc* mutant mice, which display strain-dependent phenotypic variation (Shoemaker et al. 1998); the medaka fish, where thermal stress induces sex reversal at different rates in Hd-rR versus HNI strains (Sato et al. 2005); and *Drosophila*, where mutant phenotypes of *scalloped* manifest more severely in the Oregon-R background than in the Samarkand background (Dworkin et al. 2009). Nevertheless, establishing inbred strains is very challenging in animals that formerly were rarely used for experiments, because inbreeding is inevitably accompanied by inbreeding depression, a reduction of biological fitness caused by loss of genetic diversity.

Embryos of solitary ascidians, including those of *Ciona* species, have been used for studies, including investigations in developmental biology, for over 150 years, due to the simplicity of their embryos and larvae (Satoh et al. 2003; Lemaire 2011). For example, invariant cell lineages have been almost completely revealed (Nishida 1987), and the structure of the gene regulatory network controlling cell specification is now available at single-cell resolution (Satou and Imai 2015; Satou 2020). Ascidians belong to the subphylum Tunicata, the sister group of vertebrates. Because of this unique phylogenetic position, ascidians have also been used to study the origin of vertebrates. For example, neural crest cells and neuromesodermal cells had been thought to exist only in vertebrate embryos, but we now know that ascidian embryos have cells that share their evolutionary origins with these vertebrate-specific cells (Abitua et al. 2012; Stolfi et al. 2015; Waki et al. 2015; Horie et al. 2018; Ishida and Satou 2024).

Although colonial ascidians can be maintained clonally in the laboratory (Wawrzyniak et al. 2021), no inbred strains for solitary ascidians have been available, necessitating the use of wild-caught animals. However, *Ciona intestinalis* type A (or *Ciona robusta*), which is one of the most commonly used ascidian species, has a heterozygosity rate of 1.1% (Dehal et al. 2002; Satou et al. 2012), and this high rate potentially affects reproducibility of experiments.

To resolve the potential reproducibility problem, we previously sought to establish an inbred line of this animal; we repeated self-fertilization more than twelve times, reducing heterogeneity in this line (Satou et al. 2015). However, it suffered severely from inbreeding depression, and was finally lost. Therefore, as an alternative, instead of reducing genetic variations, we have managed genetic variations. Specifically, we have kept closed colonies and monitored their genetic variation for years to ensure reproducibility of experiments and to provide traceable genetic information.

## Results

### Annually renewed, closed colonies of ascidians

Among ascidians, *Ciona intestinalis* type A is one of the species most widely used for experiments. Brunetti et al. proposed renaming this species *Ciona robusta*, with *Ciona intestinalis* type B as *Ciona intestinalis* (Brunetti et al. 2015). Although most genetic records for “*Ciona intestinalis”* in public databases, including DDBJ/EMBL/Genbank, were created without discriminating these two types of *Ciona*, the majority of them are probably for “*Ciona robusta*”, judging from nucleotide sequences and geographic locations of submitters. Animals used in the present study are *Ciona intestinalis* type A or *Ciona robusta*, and we used the genome of this species for analyses described below.

We have kept ascidians for over ten years as colonies renewed annually and as permanently closed colonies. We have monitored their genetic backgrounds and those of wild-caught animals used for establishing these colonies by sequencing their genomic DNA (Figure 1). In 2016, we began to keep two groups of ascidians as closed colonies. These two groups are derived from Onagawa (Miyagi Prefecture, Japan) and Onahama (Fukushima Prefecture, Japan), which are approximately 170 km apart. Hereafter we refer to them as Onagawa (GW)-line and Onahama (HM)-line, respectively. Every fall from 2016 to 2020, we collected ascidians in Onagawa and Onahama, and these wild-caught animals were used for establishing colonies in subsequent years (Figure 1A, C). These animals were maintained for a year with intra-population breeding. Specifically, fertilized eggs were obtained by crossing animals only with other members of the same population, and after metamorphosis we raised juveniles for two or three weeks in the laboratory. Then, we transferred these juveniles to the sea and raised them there for one to three months until they became sexually mature. This procedure was repeated several times every year. In this way, we kept two lines of colonies from 2016 to 2020, and introduced new wild animals that were caught in Onagawa and Onahama every year.

**Figure 1.**
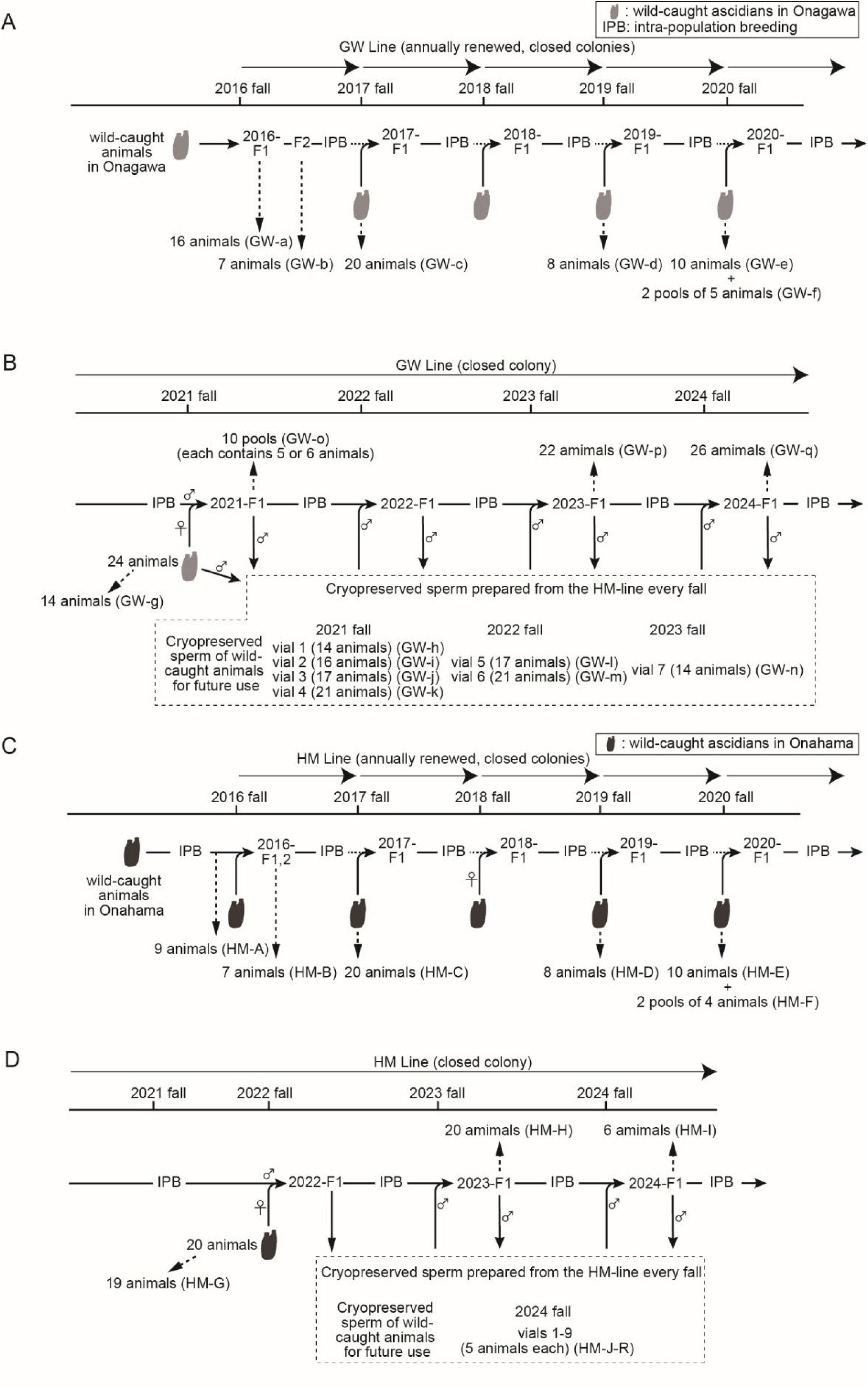
Establishment of closed colonies of ascidians. Two lines of closed colonies were established in the present study. The GW-line (A, B) began with animals collected at Onagawa, Miyagi Prefecture, Japan and the HM-line (C, D) began with animals collected at Onahama, Fukushima Prefecture, Japan. At the beginning, these two lines were re-established every year (A, C). The current two lines of closed colonies have been kept through intra-population breeding for over two or three years (B, D). Sample groups for sequencing are shown by dotted arrows. Note that sample groups GW-f, GW-o, and HM-F are pools of several animals, and the remaining samples were collected and analyzed individually. Also note that samples a, c–n, and C-G are wild-caught animals, and the remaining samples are animals we raised.

We randomly chose 7 to 20 animals from wild-caught animals that were used for crossing every year (sample groups GW-a/c/d/e/f (Figure 1A), and sample groups HM-C to F (Figure 1C)) and from the first- or second-generation animals after crossing (sample group GW-b, HM-B), and identified genetic variants using next-generation sequencing (Table S1). Although it is possible that these populations might have had undiscovered genetic variants, we identified 6,191,519 and 6,277,792 single-nucleotide substitutions for the GW- and HM-lines, respectively, which correspond to approximately 5% of the total length of the genome. As we show below, wild animals caught in Onagawa and Onahama have distinct sets of genetic variants, and variants these animals had did not drastically change during the five years of our study.

### Closed colonies of ascidians

To obtain a more genetically stable, closed colony, we caught 24 animals in Onagawa in the fall of 2021. Their eggs were crossed with sperm obtained from the GW-line that we had kept as an annual closed colony since 2020. We obtained sufficient amounts of DNA from 14 of these 24 animals, and analyzed their genomic sequences (sample groups GW-g; Figure 1B). Because we failed to obtain enough sperm from the remaining 10 animals, we also analyzed 10 pools of 5 to 6 first-generation animals after the crossing (sample groups GW-o; Figure 1B).

To keep the colony for a long time, we cryopreserved sperm of the first-generation animals. This cryopreserved sperm was used in the fall of 2022 for recovery of genetic variation, thereby preventing inbreeding depression. From the first-generation animals after this cross, we again cryopreserved sperm for future use; these cryopreserved sperm will be used to recover genetic variations when heterozygosity rates of the colonies become reduced. We repeated this step twice more in 2023 and 2024 (Figure 1B).

To monitor heterozygosity rates of these colonies, we randomly selected 22 and 26 animals in 2023 and 2024 from this population, and performed DNA-sequencing (sample groups GW-p and q; Figure 1B), because we intend not to reduce genetic diversity of these colonies. We calculated heterozygosity rates for each individual using Bcftools (Li 2011), and found that genetic diversity in the population had gradually decreased despite back-crossing (Figure 2A; Table S2). Note that heterozygosity rates estimated in the present study were lower even in wild-caught animals (sample groups GW-e and GW-g) than the heterozygosity rates estimated by different methods (1.1 to 1.2%) (Dehal et al. 2002; Satou et al. 2012). However, because the purpose of this analysis was to monitor population genetic diversity not to be reduced, we did not determine which estimates were more accurate.

**Figure 2.**
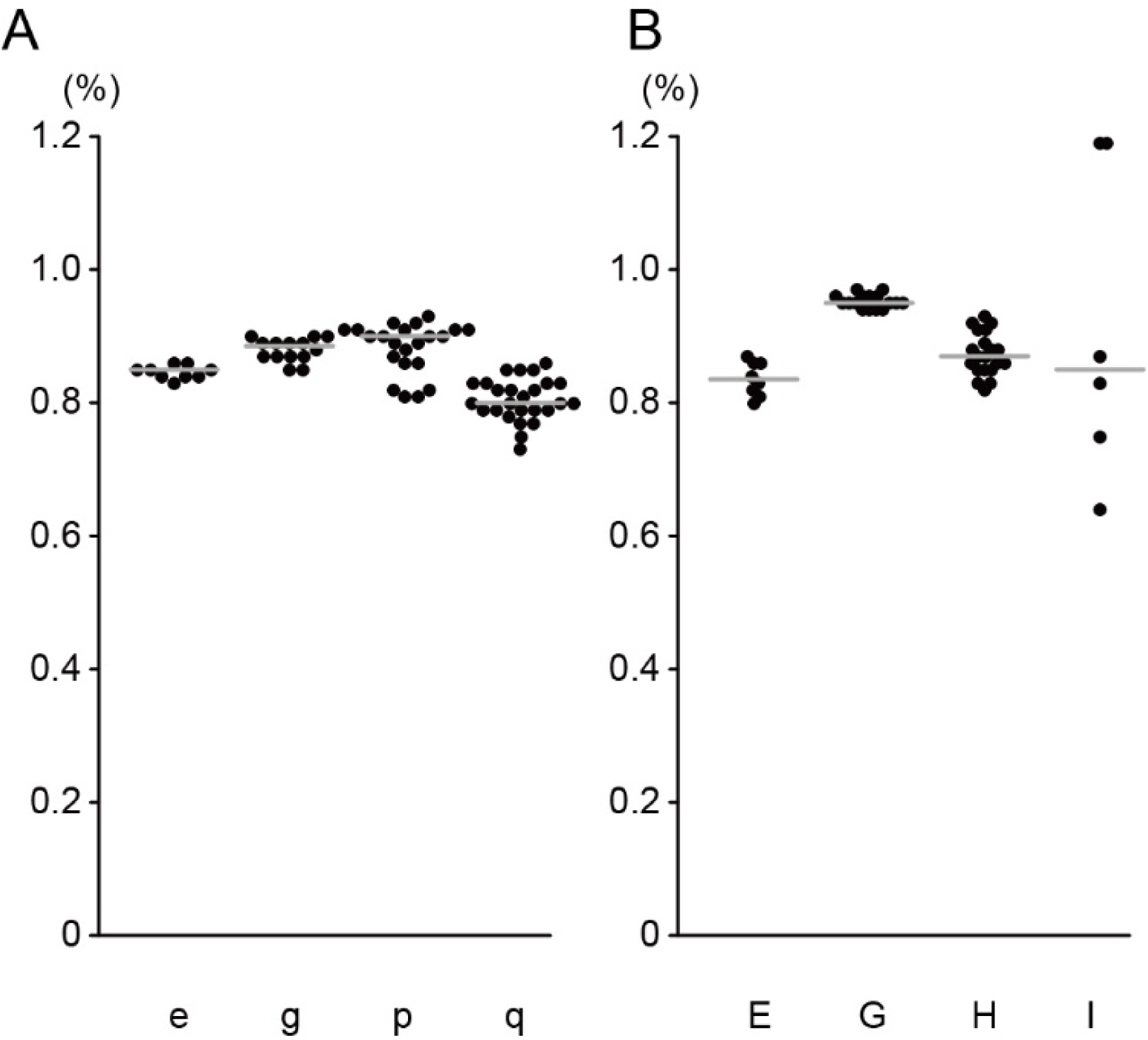
Monitoring heterozygosity rates in the permanently closed colonies to maintain genetic variations in the GW- and HM-lines. Heterozygosity rates of individuals of (A) GW- and (B) HM-lines per sample group are shown with dots. Median values are shown with gray lines. Note that samples, sequence depths of which did not exceed 20x, are not included in the graph. See all data in Table S2.

To overcome potential inbreeding depression in the future, we also prepared seven pools of cryopreserved sperm derived from animals caught in Onagawa in the fall of 2021, 2022, and 2023 (sample groups GW-h to n; Figure 1B). Genetic variation of these animals was individually analyzed (Table S1).

Similarly, in the fall of 2022, we collected 20 animals in Onahama, and crossed these animals with the HM-line (Figure 1D). We also analyzed their genetic variation from aliquots of sperm of 19 of these 20 animals. We cryopreserved sperm from first generation animals after the crossing in 2022. In the fall of 2023 and 2024, we back-crossed HM-line animals with sperm that had been cryopreserved one year earlier. After these back-crosses in 2023 and 2024, we randomly chose 20 and 6 animals (sample groups HM-H and I; Figure 1D) to monitor their heterozygosity rates. As in the case of the GW-line, the median heterozygosity rate of the HM-line decreased in 2024, although the deviation was large (Figure 2B; Table S2). We prepared 9 pools of cryopreserved sperm, each of which contained 5 individuals, to recover genetic diversity in the future (sample groups HM-J-R; Figure 1D).

### Quantitative analyses of genetic variation

To quantitatively evaluate genetic variation of animals used for establishing the above colonies, we analyzed sequencing results of sample groups GW-a–e, g–n, and HM-A–E, G, J–R, because we collected and determined genomic sequences individually for these sample groups. As controls, we determined nucleotide variation of 17 animals that were caught in Okayama (approximately 780 km from Onagawa, and 660 km apart from Onahama) in the spring and fall of 2017 (sample groups YM-V and U).

A multi-dimensional analysis for variation and hierarchical clustering indicated that GW animals and HM animals largely constitute two different groups, and these populations are distinct from the Okayama population (Figure 3AB). In particular, these analyses showed that genetic variants of animals caught at Onagawa in and after 2020 (GW-g–n) are similar, and genetic variants of animals caught at Onahama in and after 2020 (HM-G, J–R) are also similar, with a few exceptions, while genetic variants are distinct between the GW- and HM-populations.

**Figure 3.**
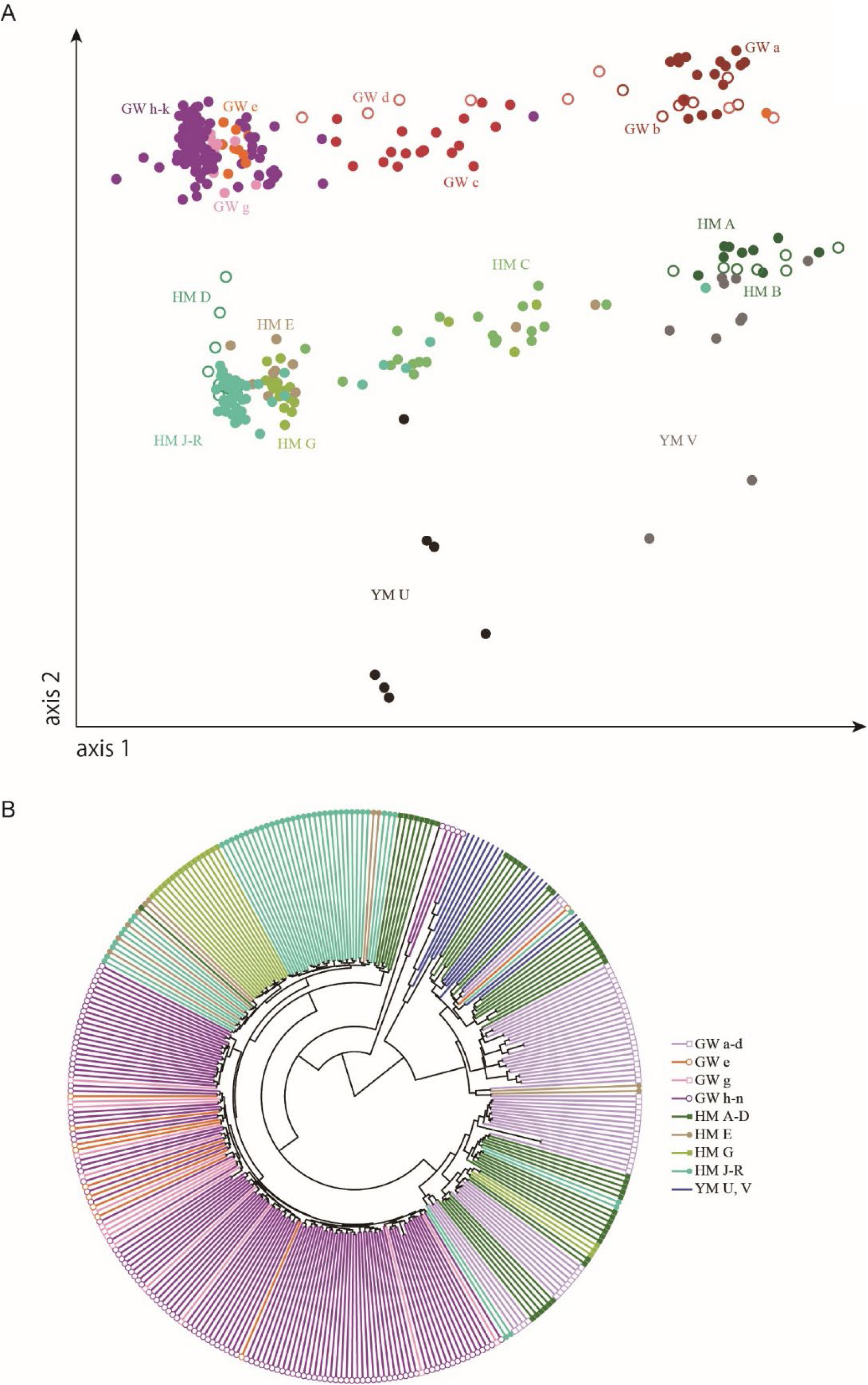
Genetic variation of ascidians derived from three locations. (A) A multi-dimensional scaling plot for sample groups GW-a–e/g–k, HM-A-E/G/J–R, and YM-U/V. (B) A tree constructed with the complete-linkage clustering method. Each branch is colored according to the legend in the panel.

All animals except animals in groups GW-b, HM-A, and HM-B were wild-caught animals. In other words, wild populations in Onagawa and Onahama have maintained distinct genetic variants, although both locations are linked directly to the Pacific Ocean, and only 170-km apart. At the same time, our data show that genetic variants in each population have changed gradually (Figure 3A).

Next, we compared genetic variation in these groups. For this purpose, we also used nucleotide sequences obtained from genomic libraries, each of which was derived from multiple individuals (GW-f, o; HM-F), as well as libraries derived from single individuals, as described above. In addition, we obtained nucleotide sequences from four libraries, each of which was made from five individuals caught in Chiba, which is approximately 340 km, 170 km, 550 km from Onagawa, Onahama, and Okayama (sample groups CH-Z). A Venn diagram (Figure 4A) indicates that these four populations have distinct genetic backgrounds. At the same time, because 218,100 (=175,913+35,785+6,402) genetic variations were found only in control wild-animal samples (YM-U, YM-V, and CH-Z), it is highly likely that many uncharacterized genetic variations are present in wild-animals.

**Figure 4.**
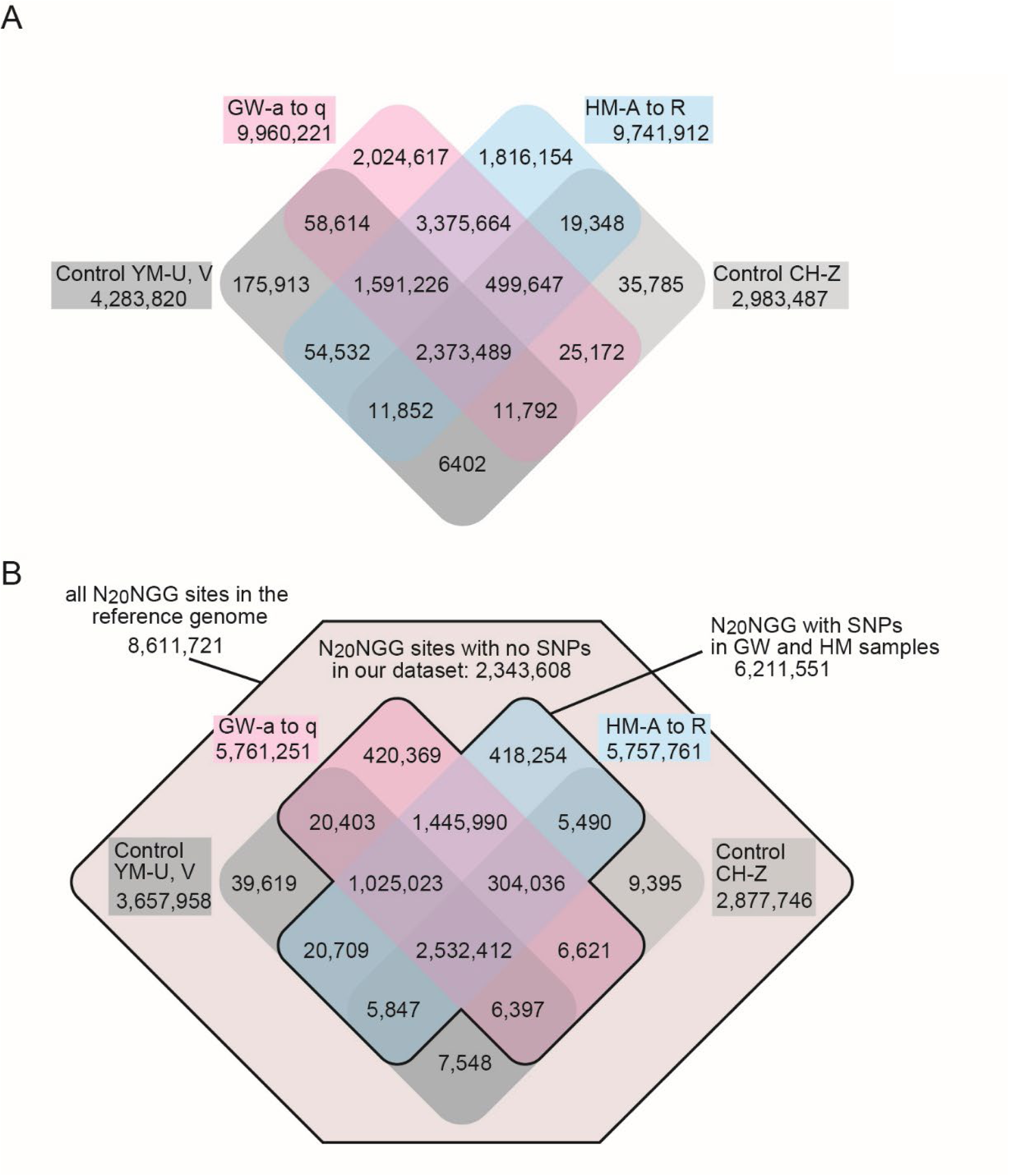
Venn-diagrams showing overlaps of variants among four populations. (A) Numbers of variants are shown. (B) Numbers of “N_20_NGG” motifs in the reference genome and those containing SNPs in the GW- and HM-lines and controls. While SNPs found in controls indicate that wild-animals have uncharacterized SNPs, all genetic variants have been characterized in the GW- and HM-lines.

Finally, to demonstrate the potential of the GW- and HM-lines to enhance experimental reproducibility, we examined SNPs located within “N_20_NGG” motifs, which serve as guide RNA and PAM sequences for CRISPR/Cas9 genome editing (Doudna and Charpentier 2014). We identified 8,611,721 N_20_NGG motifs in the reference genome; of these, 6,211,551 (72%) contained SNPs in the GW- and/or HM-samples (Figure 4B). Excluding these variants yields 2,850,470 (=8,611,721-5,761,251) and 2,853,960 (=8,611,721-5,757,761) usable N_20_NGG sites for GW-line and HM-line, respectively. Because the genetic variations of the GW- and HM-lines have been fully characterized, high reproducibility is ensured by selecting CRISPR/Cas9 targets that avoid these known SNPs of the GW- and HM-lines and by using GW- and HM-line animals. In contrast, wild populations may have uncharacterized SNPs within these N_20_NGG sites that are usable sites in GW- and HM-lines. This was confirmed by the observation of 56,562 (=39,619+7,548+9,395) SNPs in the control YM-and CH-animals that were absent in the GW- and HM-lines, which highlights a risk of reducing editing efficiency and experimental reproducibility in use of wild animals.

## Discussion

In the present study, we found that four populations are genetically distinct. These variants are easily accessible in a genome browser common to the *Ciona* research community (https://ghost.zool.kyoto-u.ac.jp/default_ht.html) (Satou et al. 2005). Researchers can increase reproducibility of experiments by designing PCR primers, antisense oligonucleotides, probes for HCR *in situ* hybridization, and guide RNAs for CRISPR-knockout in genomic regions in which no variants are found and by using GW- and HM-line animals.

We recommend that researchers use the GW- and HM-lines in their experiments and publish the dates when experimental animals they used were obtained. This will increase reproducibility and credibility of experimental results.

The cost of next-generation sequencing has been steadily decreasing, and the number of genome sequences decoded is increasing. This makes more “non-model” organisms useful for experiments. Our establishment of annual and permanently closed colonies of ascidians will provide a cost-effective and feasible way to increase and ensure reproducibility of experiments using such new experimental models.

## Materials and Methods

### Animals

Animals used to establish the GW-line were collected in the sea near the Onagawa Field Center, Graduate School of Agricultural Science, Tohoku University. Animals used to establish the HM-line were caught near the Aquamarine Fukushima Aquarium. Control animals were collected in the sea near the Ushimado Marine Institute of Okayama University, and also near the Chiba port.

### Breeding

Eggs and sperm were obtained by dissecting animals. We incubated fertilized eggs until hatching. Healthy larvae were transferred to plastic dishes at the approximate concentration of 0.16 animals / cm^2^, i.e., 10 animals per 9-cm dish. After two or three weeks, incubated animals were transferred into the sea in two locations. The first was a pier of the Field Science Education and Research Center of Kyoto University in Maizuru City, Kyoto Prefecture. The second was an aquaculture raft of the Misaki Marine Biological Station of the University of Tokyo, in Miura City, Kanagawa Prefecture.

Cryopreservation of sperm was performed according to a previous method with minor modifications (Moody et al. 1999). Sperm were collected from sperm ducts of dissected animals, taking care to minimize the amount of other fluid. Collected sperm were kept on ice until they were frozen. Sperm were mixed with 10% dimethylsulfoxide (v/v) in seawater, at a ratio of 1:4. This mixture was divided into aliquots in cryostorage vials. After insertion into cryocanes, vials were immediately immersed in liquid nitrogen.

For use, a vial including cryopreserved sperm was taken from the cane, and seawater was immediately poured directly into the vial. After gentle mixing several times with a glass pipette, thawed sperm were transferred to seawater containing unfertilized eggs for artificial fertilization.

### Genome sequencing

Genomic DNA was isolated from sperm. DNA libraries were made with a Nextera XT DNA Library Preparation Kit, a Nextera DNA Flex library preparation kit, an Illumina DNA prep kit, or an Illumina DNA PCR-Free Prep kit, and analyzed with Illumina sequencers. All sequences are deposited in the DRA/SRA database under accession numbers DRR725991–DRR726412. Paired-end reads were mapped to the reference genome (HT version) (Satou et al. 2019) using Bowtie2 with default options (Langmead and Salzberg 2012). Obtained sam files were sorted and converted to bam-formatted files with Samtools (Li et al. 2009). The “mpileup” command of bcftools (Li 2011) was used to generate bcf files containing genotype likelihoods with the option “-Ob.” Then the “call” command of bcftools was used for calling single nucleotide variants and insertions/deletions with options “-Ob” and “-m”, and all bcf files were merged with the “merge” command of bcftools with the option “-m b.” After indexing the files, we used the “view” command of bcftools to choose only positions that include single-nucleotide variants and insertions/deletions. Then, we split multiallelic records into individual records using the “norm” command with options “-m” and “-Ou”, left-aligned insertions/deletions using the “norm” command with options “-f” and “-Ou” and the reference genome sequence, removed duplicated records using the “norm” command with options “-d both” and “-Ou”, and gave a unique name to each variant using the “annotate” command with options “-Ob”, “-x ID”, and “-I +’%CHROM:%POS:%REF:%ALT’”. To calculate numbers of overlapping variants, we used the “isec” command.

Using the obtained BCF file, we analyzed variants with Plink (version 1.90b6.21) (Chang et al. 2015). All commands used for making the MDS plot are shown in Supplemental Text 1. To estimate heterozygosity rates, we used the “stats” command of bcftools with the option “-s -”. We also estimated the number of non-duplicated reads in each library using Fastqc (www.bioinformatics.babraham.ac.uk/projects/fastqc/). Depth of non-duplicated reads was calculated assuming a genome size of 125 Mb (Table S1). To summarize heterozygosity rates, we used data for which read depths exceeded 20x.

## Data access

All sequences are deposited in the DRA/SRA database under accession numbers DRR725991–DRR726412.

## Competing interests

We declare no competing interests.

## Acknowledgements

We thank all staff members at the Maizuru Fisheries Research Station of Kyoto University, and the Misaki Marine Biological Station of the University of Tokyo, the Onagawa Field Center of Tohoku University, AquaMarine Fukushima, and the Ushimado Marine Institute of Okayama University for their cooperation in collection of animals. We also thank Ms. Chikako Imaizumi, Mr. Yoshikazu Okada, Ms. Yoko Kitagawa, Ms. Reiko Sakota, Ms. Kanako Nagai, Ms. Yuriko Murata, Ms. Michiko Hosotani, Mr. Koji Hosotani, Ms. Natsumi Maeda (Kyoto University), Ms. Atsuko Ishiwata, Ms. Reiwa Igarashi, Dr. Yohei Otomo, Ms. Michiyo Kawabata, Mr. Yoshiaki Uchida, Mr. Makoto Nozawa (the University of Tokyo), Mr. Kazuhiro Saito, and Mr. Waichiro Godo (Okayama University) for their technical assistance, and to Prof. Koji Akasaka and Prof. Toru Miura for their generous support in allowing us to use the Misaki Marine Biological Station facility.

This research was funded by the National BioResource Project of Japanese Ministry of Education, Culture, Sports, Science, and Technology / Japan Science and Technology Agency / Japan Agency for Medical Research and Development.

## Author contributions

Yutaka Satou: Writing – original draft preparation; Conceptualization; Investigation; Resources; Manabu Yoshida: Writing – original draft preparation; Conceptualization; Resources; Yasunori Sasakura : Writing – original draft preparation; Conceptualization; Shin-ichi Tokuhiro: Investigation; Reiko Yoshida: Investigation; Resources; Kazuo Inaba: Conceptualization; Resources; Kogiku Shiba: Resources; Mayuko Hamada: Resources; Nori Satoh : Investigation; Conceptualization; Aika Shibata : Resources; Hisanori Kohtsuka: Resources; Hiromi Kakizaki: Resources; Akihiro Yoshikawa: Resources; Satoe Aratake: Resources; Michio Ogasawara: Resources; Reiji Masuda: Resources; Takehiro G. Kusakabe: Resources

## Supplementary Text S1. Markdowm in the analysis using Plink and R

\# input bcf file: allvariants.bcf; this bcf file includes animals in sample groups GW-a–e/g–k, HM-A-E/G/J–R, and YM-U/V.
$ plink --bcf allvariants.bcf --keep-allele-order -vcf-idspace-to _ --const-fid --allow-extra-chr 0 --id-delim - --make-bed --out AllSample.1
\# 11537155 variants and 330 people pass filters and QC

\# exclude variants that are not called in 2% or more animals
$ plink --bfile AllSample.1 --geno 0.02 --make-bed --out AllSample.2
\# 6515265 variants and 330 people pass filters and QC

\# exclude animals in which 60% or more variant positions failed to be called
$ plink --bfile AllSample.2 --mind 0.6 --make-bed --out AllSample.3
\# 6515265 variants and 330 people pass filters and QC; No animals were excluded.

\# exclude low minor allele frequency variants (5% or less)
$ plink --bfile AllSample.3 --maf 0.05 --make-bed --out AllSample.4
\# 2608683 variants and 330 people pass filters and QC

\# keep only variants in Hardy-Weinberg equilibrium (<1e-6)
$ plink --bfile AllSample.4 --make-bed --out AllSample.5 --hwe include-nonctrl 1e-6
\# 2237171 variants and 330 people pass filters and QC

#Clustering
$ plink --bfile AllSample.5 --cluster --mds-plot 2 --out AllSample.6-mds
$ plink --bfile AllSample.5 --distance square allele-ct --out AllSample.6-mat
$ R
> mds <- read.table(" AllSample.6-mds.mds", header=T)
> plot(mds$C1, mds$C2)
> data <- read.table(" AllSample.6-mat.dist")
> data_id <- read.table(" AllSample.6-mat.dist.id")
> row.names(dat)<-c(data_id[,s])
> hc <- hclust(dist(data))
> library(ape)
> plot(as.phylo(hc), type="fan")
> q()

---

**Table S1.** Basic statistics of sequencing analyses.

| Origin*1 | Sample Group | IDs | Number of reads | Number of nucleotides | Percentage of non-duplicated reads*2 | Number of nucleotides of deduplicated reads | Depth*3 |
| --- | --- | --- | --- | --- | --- | --- | --- |
| GW | a | 1 | 13,137,560 | 2,684,385,534 | 88% | 403,929,950 | 3.3 |
| GW | a | 2 | 12,705,096 | 2,861,378,956 | 88% | 389,844,603 | 3.2 |
| GW | a | 3 | 11,141,076 | 2,466,290,723 | 88% | 341,493,636 | 2.8 |
| GW | a | 4 | 11,171,878 | 2,368,370,896 | 87% | 339,476,373 | 2.8 |
| GW | a | 5 | 13,168,826 | 2,873,011,117 | 87% | 401,324,489 | 3.3 |
| GW | a | 6 | 12,278,108 | 2,618,167,564 | 86% | 368,818,304 | 3.0 |
| GW | a | 7 | 14,005,002 | 3,049,597,255 | 88% | 429,418,569 | 3.5 |
| GW | a | 8 | 11,057,406 | 2,467,707,915 | 88% | 339,196,099 | 2.8 |
| GW | a | 9 | 11,607,096 | 2,659,638,665 | 87% | 355,213,670 | 2.9 |
| GW | a | 10 | 12,294,446 | 2,647,576,697 | 87% | 373,246,128 | 3.0 |
| GW | a | 11 | 11,316,272 | 2,590,421,551 | 87% | 343,303,983 | 2.8 |
| GW | a | 12 | 12,622,256 | 2,635,796,047 | 86% | 379,272,119 | 3.1 |
| GW | a | 13 | 14,721,804 | 3,309,039,577 | 87% | 445,942,340 | 3.6 |
| GW | a | 14 | 12,764,494 | 2,832,174,649 | 86% | 384,965,623 | 3.1 |
| GW | a | 15 | 12,623,210 | 2,719,579,586 | 87% | 383,681,814 | 3.1 |
| GW | a | 16 | 11,961,064 | 2,708,269,539 | 90% | 375,382,378 | 3.1 |
| GW | b | 1 | 11,111,538 | 2,559,853,831 | 88% | 342,430,270 | 2.8 |
| GW | b | 2 | 12,857,074 | 2,731,858,552 | 88% | 394,057,302 | 3.2 |
| GW | b | 3 | 9,648,108 | 2,068,616,334 | 90% | 305,460,874 | 2.5 |
| GW | b | 4 | 14,435,516 | 2,307,200,614 | 90% | 456,478,958 | 3.7 |
| GW | b | 5 | 12,620,036 | 2,576,240,998 | 89% | 394,606,048 | 3.2 |
| GW | b | 6 | 12,498,392 | 2,567,785,590 | 84% | 368,747,594 | 3.0 |
| GW | b | 7 | 13,935,026 | 3,089,826,887 | 88% | 427,507,162 | 3.5 |
| GW | c | 1 | 14,343,418 | 2,948,551,021 | 86% | 432,037,879 | 3.5 |
| GW | c | 2 | 12,532,600 | 2,598,816,118 | 86% | 378,796,092 | 3.1 |
| GW | c | 3 | 14,493,088 | 3,237,827,688 | 86% | 436,603,438 | 3.5 |
| GW | c | 4 | 12,797,564 | 2,459,735,426 | 86% | 383,105,767 | 3.1 |
| GW | c | 5 | 14,887,314 | 3,206,704,254 | 85% | 445,198,305 | 3.6 |
| GW | c | 6 | 14,253,314 | 2,951,496,547 | 88% | 437,422,107 | 3.6 |
| GW | c | 7 | 11,714,562 | 2,452,033,721 | 88% | 360,138,082 | 2.9 |
| GW | c | 8 | 12,462,386 | 2,660,789,423 | 84% | 365,638,587 | 3.0 |
| GW | c | 9 | 8,899,124 | 1,744,451,781 | 88% | 275,272,419 | 2.2 |
| GW | c | 10 | 9,079,150 | 1,897,896,864 | 87% | 275,162,889 | 2.2 |
| GW | c | 11 | 10,881,292 | 2,225,501,653 | 88% | 334,450,756 | 2.7 |
| GW | c | 12 | 11,427,804 | 2,132,039,892 | 88% | 350,042,804 | 2.8 |
| GW | c | 13 | 14,899,798 | 3,261,129,849 | 81% | 423,535,275 | 3.4 |
| GW | c | 14 | 12,058,706 | 2,507,787,195 | 88% | 372,699,733 | 3.0 |
| GW | c | 15 | 15,882,930 | 3,452,450,771 | 85% | 474,128,753 | 3.9 |
| GW | c | 16 | 14,931,018 | 3,114,464,760 | 86% | 448,569,738 | 3.6 |
| GW | c | 17 | 11,017,044 | 2,244,633,208 | 86% | 331,113,295 | 2.7 |
| GW | c | 18 | 11,382,094 | 2,480,741,673 | 87% | 347,171,127 | 2.8 |
| GW | c | 19 | 14,498,348 | 3,101,737,423 | 88% | 444,981,675 | 3.6 |
| GW | c | 20 | 15,166,668 | 3,088,224,025 | 88% | 466,476,901 | 3.8 |
| GW | d | 1 | 39,832,514 | 5,974,877,100 | 34% | 2,059,666,914 | 16.7 |
| GW | d | 2 | 24,438,530 | 3,665,779,500 | 37% | 1,340,543,504 | 10.9 |
| GW | d | 3 | 26,318,404 | 3,947,760,600 | 55% | 2,171,552,679 | 17.7 |
| GW | d | 4 | 24,088,432 | 3,613,264,800 | 37% | 1,319,397,576 | 10.7 |
| GW | d | 5 | 29,303,920 | 4,395,588,000 | 50% | 2,194,431,261 | 17.8 |
| GW | d | 6 | 36,535,870 | 5,480,380,500 | 53% | 2,922,129,555 | 23.8 |
| GW | d | 7 | 28,047,996 | 4,207,199,400 | 26% | 1,114,591,102 | 9.1 |
| GW | d | 8 | 44,453,012 | 6,667,951,800 | 48% | 3,174,121,397 | 25.8 |
| GW | e | 1 | 42,137,046 | 6,320,556,900 | 60% | 3,782,997,220 | 30.8 |
| GW | e | 2 | 38,245,790 | 5,736,868,500 | 22% | 1,286,238,760 | 10.5 |
| GW | e | 3 | 41,722,006 | 6,258,300,900 | 59% | 3,706,905,745 | 30.1 |
| GW | e | 4 | 41,075,932 | 6,161,389,800 | 59% | 3,646,442,792 | 29.6 |
| GW | e | 5 | 39,002,270 | 5,850,340,500 | 62% | 3,612,375,179 | 29.4 |
| GW | e | 6 | 36,550,008 | 5,482,501,200 | 61% | 3,339,100,975 | 27.1 |
| GW | e | 7 | 45,872,012 | 6,880,801,800 | 60% | 4,111,775,673 | 33.4 |
| GW | e | 8 | 34,251,386 | 5,137,707,900 | 63% | 3,223,085,077 | 26.2 |
| GW | e | 9 | 42,069,542 | 6,310,431,300 | 62% | 3,889,045,971 | 31.6 |
| GW | e | 10 | 34,230,748 | 5,134,612,200 | 62% | 3,182,342,165 | 25.9 |
| GW | f | 1 | 40,094,594 | 6,014,189,100 | 55% | 3,285,543,513 | 26.7 |
| GW | f | 2 | 42,714,016 | 6,407,102,400 | 63% | 4,052,858,683 | 33.0 |
| GW | g | 1 | 55,243,958 | 8,286,593,700 | 59% | 4,901,552,319 | 39.9 |
| GW | g | 2 | 54,812,432 | 8,221,864,800 | 59% | 4,862,742,926 | 39.5 |
| GW | g | 3 | 66,208,810 | 9,931,321,500 | 55% | 5,506,341,609 | 44.8 |
| GW | g | 4 | 65,846,480 | 9,876,972,000 | 57% | 5,654,594,406 | 46.0 |
| GW | g | 5 | 51,170,072 | 7,675,510,800 | 58% | 4,466,482,952 | 36.3 |
| GW | g | 6 | 63,115,920 | 9,467,388,000 | 58% | 5,467,568,636 | 44.5 |
| GW | g | 7 | 57,908,958 | 8,686,343,700 | 59% | 5,124,949,207 | 41.7 |
| GW | g | 8 | 56,943,734 | 8,541,560,100 | 59% | 5,000,634,254 | 40.7 |
| GW | g | 9 | 60,373,436 | 9,056,015,400 | 57% | 5,169,091,427 | 42.0 |
| GW | g | 10 | 61,150,390 | 9,172,558,500 | 57% | 5,238,146,059 | 42.6 |
| GW | g | 11 | 61,067,146 | 9,160,071,900 | 57% | 5,254,700,879 | 42.7 |
| GW | g | 12 | 48,382,566 | 7,257,384,900 | 59% | 4,291,314,862 | 34.9 |
| GW | g | 13 | 46,692,696 | 7,003,904,400 | 59% | 4,138,789,562 | 33.6 |
| GW | g | 14 | 57,991,464 | 8,698,719,600 | 59% | 5,107,162,100 | 41.5 |
| GW | h | 1 | 67,262,148 | 10,156,584,348 | 79% | 8,016,276,219 | 65.2 |
| GW | h | 2 | 57,466,962 | 8,677,511,262 | 75% | 6,530,307,855 | 53.1 |
| GW | h | 3 | 26,280,448 | 3,942,067,200 | 60% | 2,373,067,420 | 19.3 |
| GW | h | 4 | 60,309,150 | 9,106,681,650 | 80% | 7,330,852,721 | 59.6 |
| GW | h | 5 | 64,913,662 | 9,801,962,962 | 80% | 7,818,352,707 | 63.6 |
| GW | h | 6 | 69,844,454 | 10,546,512,554 | 76% | 8,037,077,135 | 65.3 |
| GW | h | 7 | 67,774,018 | 10,233,876,718 | 67% | 6,873,076,576 | 55.9 |
| GW | h | 8 | 70,331,174 | 10,620,007,274 | 73% | 7,774,637,942 | 63.2 |
| GW | h | 9 | 73,650,548 | 11,121,232,748 | 70% | 7,776,299,344 | 63.2 |
| GW | h | 10 | 75,808,558 | 11,447,092,258 | 78% | 8,970,547,325 | 72.9 |
| GW | h | 11 | 64,319,808 | 9,712,291,008 | 72% | 7,000,043,146 | 56.9 |
| GW | h | 12 | 80,878,812 | 12,212,700,612 | 72% | 8,807,743,571 | 71.6 |
| GW | h | 13 | 76,802,058 | 11,597,110,758 | 76% | 8,772,315,379 | 71.3 |
| GW | h | 14 | 78,566,688 | 11,863,569,888 | 69% | 8,211,604,690 | 66.8 |
| GW | i | 1 | 67,092,834 | 10,131,017,934 | 75% | 7,648,448,202 | 62.2 |
| GW | i | 2 | 74,206,872 | 11,205,237,672 | 71% | 7,927,422,203 | 64.5 |
| GW | i | 3 | 70,751,444 | 10,683,468,044 | 75% | 8,046,065,508 | 65.4 |
| GW | i | 4 | 76,471,536 | 11,547,201,936 | 72% | 8,337,980,336 | 67.8 |
| GW | i | 5 | 70,653,702 | 10,668,709,002 | 74% | 7,947,331,924 | 64.6 |
| GW | i | 6 | 76,855,240 | 11,605,141,240 | 74% | 8,609,274,116 | 70.0 |
| GW | i | 7 | 73,077,448 | 11,034,694,648 | 73% | 8,038,845,519 | 65.4 |
| GW | i | 8 | 67,810,170 | 10,239,335,670 | 73% | 7,456,765,638 | 60.6 |
| GW | i | 9 | 65,791,046 | 9,934,447,946 | 75% | 7,421,212,169 | 60.3 |
| GW | i | 10 | 81,529,928 | 12,311,019,128 | 72% | 8,828,970,731 | 71.8 |
| GW | i | 11 | 66,357,902 | 10,020,043,202 | 74% | 7,420,402,037 | 60.3 |
| GW | i | 12 | 78,831,598 | 11,903,571,298 | 70% | 8,385,337,210 | 68.2 |
| GW | i | 13 | 75,786,164 | 11,443,710,764 | 75% | 8,578,874,657 | 69.7 |
| GW | i | 14 | 83,120,382 | 12,551,177,682 | 70% | 8,759,007,816 | 71.2 |
| GW | i | 15 | 74,412,850 | 11,236,340,350 | 73% | 8,190,934,854 | 66.6 |
| GW | i | 16 | 119,421,776 | 18,032,688,176 | 73% | 13,148,686,085 | 106.9 |
| GW | j | 1 | 77,731,982 | 11,737,529,282 | 71% | 8,388,289,900 | 68.2 |
| GW | j | 2 | 78,732,258 | 11,888,570,958 | 72% | 8,596,302,141 | 69.9 |
| GW | j | 3 | 33,334,310 | 5,008,836,192 | 27% | 1,345,952,026 | 10.9 |
| GW | j | 4 | 70,795,214 | 10,690,077,314 | 71% | 7,634,399,025 | 62.1 |
| GW | j | 5 | 77,453,562 | 11,695,487,862 | 73% | 8,588,465,822 | 69.8 |
| GW | j | 6 | 74,112,232 | 11,190,947,032 | 70% | 7,872,585,607 | 64.0 |
| GW | j | 7 | 68,529,370 | 10,347,934,870 | 68% | 7,057,098,283 | 57.4 |
| GW | j | 8 | 71,314,160 | 10,768,438,160 | 69% | 7,443,008,464 | 60.5 |
| GW | j | 9 | 70,662,554 | 10,670,045,654 | 74% | 7,843,448,795 | 63.8 |
| GW | j | 10 | 24,914,130 | 1,868,559,750 | 60% | 2,254,026,895 | 18.3 |
| GW | j | 11 | 68,285,044 | 10,311,041,644 | 71% | 7,360,897,470 | 59.8 |
| GW | j | 12 | 73,560,558 | 11,107,644,258 | 72% | 8,005,951,311 | 65.1 |
| GW | j | 13 | 74,409,020 | 11,235,762,020 | 74% | 8,310,397,678 | 67.6 |
| GW | j | 14 | 81,800,082 | 12,351,812,382 | 73% | 8,989,491,245 | 73.1 |
| GW | j | 15 | 81,877,728 | 12,363,536,928 | 71% | 8,759,855,838 | 71.2 |
| GW | j | 16 | 78,074,370 | 11,789,229,870 | 72% | 8,431,273,595 | 68.5 |
| GW | j | 17 | 82,663,288 | 12,482,156,488 | 72% | 8,938,320,285 | 72.7 |
| GW | k | 1 | 21,568,133 | 6,470,000,000 | 62% | 4,033,041,987 | 32.8 |
| GW | k | 2 | 28,707,673 | 8,612,000,000 | 58% | 4,993,665,165 | 40.6 |
| GW | k | 3 | 21,556,880 | 6,467,000,000 | 64% | 4,139,251,195 | 33.7 |
| GW | k | 4 | 34,648,235 | 10,394,000,000 | 55% | 5,763,082,447 | 46.9 |
| GW | k | 5 | 17,092,015 | 5,128,000,000 | 66% | 3,377,961,616 | 27.5 |
| GW | k | 6 | 19,947,112 | 5,984,000,000 | 64% | 3,816,060,509 | 31.0 |
| GW | k | 7 | 18,583,226 | 5,575,000,000 | 58% | 3,245,845,152 | 26.4 |
| GW | k | 8 | 20,280,480 | 6,084,000,000 | 58% | 3,535,809,262 | 28.7 |
| GW | k | 9 | 19,349,020 | 5,805,000,000 | 64% | 3,700,305,239 | 30.1 |
| GW | k | 10 | 14,954,985 | 4,486,000,000 | 66% | 2,979,246,383 | 24.2 |
| GW | k | 11 | 19,394,528 | 5,818,000,000 | 62% | 3,599,046,871 | 29.3 |
| GW | k | 12 | 6,544,884 | 1,963,000,000 | 67% | 1,313,783,363 | 10.7 |
| GW | k | 13 | 34,523,816 | 10,357,000,000 | 59% | 6,154,932,916 | 50.0 |
| GW | k | 14 | 21,654,040 | 6,496,000,000 | 65% | 4,220,965,615 | 34.3 |
| GW | k | 15 | 21,066,960 | 6,320,000,000 | 65% | 4,098,856,366 | 33.3 |
| GW | k | 16 | 12,087,316 | 3,626,000,000 | 67% | 2,436,872,164 | 19.8 |
| GW | k | 17 | 16,771,912 | 5,032,000,000 | 66% | 3,343,642,423 | 27.2 |
| GW | k | 18 | 16,091,391 | 4,827,000,000 | 69% | 3,312,898,101 | 26.9 |
| GW | k | 19 | 20,744,381 | 6,223,000,000 | 63% | 3,905,647,162 | 31.8 |
| GW | k | 20 | 21,341,546 | 6,402,000,000 | 64% | 4,105,776,809 | 33.4 |
| GW | k | 21 | 21,912,295 | 6,574,000,000 | 67% | 4,380,203,102 | 35.6 |
| GW | l | 1 | 80,261,846 | 12,119,538,746 | 74% | 8,938,489,578 | 72.7 |
| GW | l | 2 | 92,531,912 | 13,972,318,712 | 73% | 10,194,766,751 | 82.9 |
| GW | l | 3 | 81,963,620 | 12,376,506,620 | 72% | 8,902,599,298 | 72.4 |
| GW | l | 4 | 72,390,226 | 10,930,924,126 | 74% | 8,102,683,702 | 65.9 |
| GW | l | 5 | 78,211,694 | 11,809,965,794 | 73% | 8,583,583,277 | 69.8 |
| GW | l | 6 | 75,239,522 | 11,361,167,822 | 73% | 8,314,513,625 | 67.6 |
| GW | l | 7 | 76,760,572 | 11,590,846,372 | 74% | 8,521,951,443 | 69.3 |
| GW | l | 8 | 77,907,488 | 11,764,030,688 | 72% | 8,489,800,429 | 69.0 |
| GW | l | 9 | 80,031,576 | 12,084,767,976 | 72% | 8,713,683,064 | 70.8 |
| GW | l | 10 | 72,761,218 | 10,986,943,918 | 74% | 8,184,170,431 | 66.5 |
| GW | l | 11 | 76,742,270 | 11,588,082,770 | 74% | 8,629,727,685 | 70.2 |
| GW | l | 12 | 70,161,072 | 10,594,321,872 | 73% | 7,728,037,076 | 62.8 |
| GW | l | 13 | 69,607,814 | 10,510,779,914 | 69% | 7,281,808,018 | 59.2 |
| GW | l | 14 | 76,685,486 | 11,579,508,386 | 76% | 8,762,634,859 | 71.2 |
| GW | l | 15 | 74,938,378 | 11,315,695,078 | 71% | 8,012,251,000 | 65.1 |
| GW | l | 16 | 70,303,234 | 10,615,788,334 | 75% | 7,959,280,598 | 64.7 |
| GW | l | 17 | 109,522,520 | 16,537,900,520 | 72% | 11,827,646,873 | 96.2 |
| GW | m | 1 | 83,835,326 | 12,659,134,226 | 75% | 9,521,274,610 | 77.4 |
| GW | m | 2 | 80,721,068 | 12,188,881,268 | 72% | 8,802,900,933 | 71.6 |
| GW | m | 3 | 67,441,408 | 10,183,652,608 | 75% | 7,670,682,405 | 62.4 |
| GW | m | 4 | 84,350,584 | 12,736,938,184 | 73% | 9,293,231,032 | 75.6 |
| GW | m | 5 | 54,790,044 | 8,273,296,644 | 75% | 6,187,876,813 | 50.3 |
| GW | m | 6 | 83,775,282 | 12,650,067,582 | 75% | 9,462,386,847 | 76.9 |
| GW | m | 7 | 80,542,378 | 12,161,899,078 | 75% | 9,079,759,724 | 73.8 |
| GW | m | 8 | 73,984,550 | 11,171,667,050 | 75% | 8,374,567,467 | 68.1 |
| GW | m | 9 | 75,661,104 | 11,424,826,704 | 76% | 8,633,576,811 | 70.2 |
| GW | m | 10 | 86,797,354 | 13,106,400,454 | 74% | 9,656,055,922 | 78.5 |
| GW | m | 11 | 67,760,578 | 10,231,847,278 | 77% | 7,831,976,259 | 63.7 |
| GW | m | 12 | 78,225,538 | 11,812,056,238 | 74% | 8,778,558,552 | 71.4 |
| GW | m | 13 | 81,028,432 | 12,235,293,232 | 75% | 9,181,223,089 | 74.6 |
| GW | m | 14 | 80,236,164 | 12,115,660,764 | 74% | 9,012,361,531 | 73.3 |
| GW | m | 15 | 82,561,076 | 12,466,722,476 | 74% | 9,257,308,575 | 75.3 |
| GW | m | 16 | 71,390,214 | 10,779,922,314 | 75% | 8,081,648,836 | 65.7 |
| GW | m | 17 | 80,342,216 | 12,131,674,616 | 75% | 9,103,991,614 | 74.0 |
| GW | m | 18 | 75,232,916 | 11,360,170,316 | 73% | 8,294,090,873 | 67.4 |
| GW | m | 19 | 80,899,766 | 12,215,864,666 | 75% | 9,162,212,705 | 74.5 |
| GW | m | 20 | 81,433,890 | 12,296,517,390 | 72% | 8,901,316,061 | 72.4 |
| GW | m | 21 | 68,579,120 | 10,355,447,120 | 79% | 8,218,673,106 | 66.8 |
| GW | n | 1 | 77,708,182 | 11,733,935,482 | 71% | 8,281,845,813 | 67.3 |
| GW | n | 2 | 62,677,904 | 9,464,363,504 | 75% | 7,133,289,648 | 58.0 |
| GW | n | 3 | 38,057,482 | 5,746,679,782 | 50% | 2,877,684,367 | 23.4 |
| GW | n | 4 | 23,878,154 | 3,581,723,100 | 62% | 2,205,098,850 | 17.9 |
| GW | n | 5 | 81,067,134 | 12,241,137,234 | 75% | 9,142,177,187 | 74.3 |
| GW | n | 6 | 74,661,658 | 11,273,910,358 | 73% | 8,263,712,817 | 67.2 |
| GW | n | 7 | 73,554,930 | 11,106,794,430 | 75% | 8,284,789,231 | 67.4 |
| GW | n | 8 | 75,857,448 | 11,454,474,648 | 72% | 8,244,897,585 | 67.0 |
| GW | n | 9 | 81,241,038 | 12,267,396,738 | 75% | 9,158,356,024 | 74.5 |
| GW | n | 10 | 70,502,204 | 10,645,832,804 | 75% | 7,936,273,724 | 64.5 |
| GW | n | 11 | 89,535,908 | 13,519,922,108 | 76% | 10,231,400,389 | 83.2 |
| GW | n | 12 | 78,390,354 | 11,836,943,454 | 75% | 8,883,093,388 | 72.2 |
| GW | n | 13 | 80,697,542 | 12,185,328,842 | 77% | 9,329,433,272 | 75.8 |
| GW | n | 14 | 74,226,424 | 11,208,190,024 | 76% | 8,543,593,558 | 69.5 |
| GW | o | 1 | 37,959,325 | 11,388,000,000 | 53% | 6,049,005,215 | 49.2 |
| GW | o | 2 | 46,689,237 | 14,007,000,000 | 52% | 7,314,295,032 | 59.5 |
| GW | o | 3 | 51,109,129 | 15,333,000,000 | 52% | 7,982,404,279 | 64.9 |
| GW | o | 4 | 46,321,127 | 13,896,000,000 | 53% | 7,305,496,567 | 59.4 |
| GW | o | 5 | 49,185,521 | 14,756,000,000 | 50% | 7,409,511,534 | 60.2 |
| GW | o | 6 | 45,823,633 | 13,747,000,000 | 51% | 7,065,877,655 | 57.4 |
| GW | o | 7 | 43,006,269 | 12,902,000,000 | 51% | 6,603,709,802 | 53.7 |
| GW | o | 8 | 40,356,403 | 12,107,000,000 | 45% | 5,455,885,464 | 44.4 |
| GW | o | 9 | 42,772,046 | 12,832,000,000 | 53% | 6,739,404,402 | 54.8 |
| GW | o | 10 | 39,767,122 | 11,930,000,000 | 49% | 5,797,424,439 | 47.1 |
| GW | p | 1 | 78,968,780 | 11,924,285,780 | 63% | 7,554,061,810 | 61.4 |
| GW | p | 2 | 63,521,062 | 9,591,680,362 | 78% | 7,528,923,825 | 61.2 |
| GW | p | 3 | 71,491,836 | 10,795,267,236 | 78% | 8,391,979,950 | 68.2 |
| GW | p | 4 | 68,026,526 | 10,272,005,426 | 76% | 7,834,553,334 | 63.7 |
| GW | p | 5 | 66,542,828 | 10,047,967,028 | 76% | 7,618,099,272 | 61.9 |
| GW | p | 6 | 79,835,222 | 12,055,118,522 | 78% | 9,368,315,220 | 76.2 |
| GW | p | 7 | 59,453,520 | 8,977,481,520 | 80% | 7,150,268,895 | 58.1 |
| GW | p | 8 | 66,907,676 | 10,103,059,076 | 77% | 7,756,529,168 | 63.1 |
| GW | p | 9 | 73,692,094 | 11,127,506,194 | 77% | 8,600,427,317 | 69.9 |
| GW | p | 10 | 76,821,260 | 11,600,010,260 | 75% | 8,663,420,769 | 70.4 |
| GW | p | 11 | 76,607,956 | 11,567,801,356 | 75% | 8,636,495,220 | 70.2 |
| GW | p | 12 | 87,106,936 | 13,153,147,336 | 74% | 9,784,548,186 | 79.5 |
| GW | p | 13 | 93,360,364 | 14,097,414,964 | 75% | 10,503,551,609 | 85.4 |
| GW | p | 14 | 76,702,836 | 11,582,128,236 | 78% | 9,047,912,825 | 73.6 |
| GW | p | 15 | 73,804,938 | 11,144,545,638 | 78% | 8,713,204,753 | 70.8 |
| GW | p | 16 | 67,589,390 | 10,205,997,890 | 77% | 7,815,822,645 | 63.5 |
| GW | p | 17 | 63,186,200 | 9,541,116,200 | 81% | 7,688,317,647 | 62.5 |
| GW | p | 18 | 69,523,282 | 10,498,015,582 | 79% | 8,271,562,496 | 67.2 |
| GW | p | 19 | 65,693,460 | 9,919,712,460 | 81% | 8,022,428,720 | 65.2 |
| GW | p | 20 | 75,004,554 | 11,325,687,654 | 78% | 8,783,198,021 | 71.4 |
| GW | p | 21 | 131,915,650 | 19,919,263,150 | 68% | 13,588,816,130 | 110.5 |
| GW | p | 22 | 79,645,810 | 12,026,517,310 | 72% | 8,675,677,366 | 70.5 |
| GW | q | 1 | 34,650,890 | 5,197,633,500 | 62% | 3,226,158,253 | 26.2 |
| GW | q | 2 | 34,028,834 | 5,104,325,100 | 61% | 3,108,196,107 | 25.3 |
| GW | q | 3 | 41,452,618 | 6,217,892,700 | 58% | 3,602,990,768 | 29.3 |
| GW | q | 4 | 35,629,874 | 5,344,481,100 | 61% | 3,265,422,671 | 26.5 |
| GW | q | 5 | 34,013,916 | 5,102,087,400 | 63% | 3,212,368,158 | 26.1 |
| GW | q | 6 | 36,898,994 | 5,534,849,100 | 60% | 3,335,265,906 | 27.1 |
| GW | q | 7 | 37,253,592 | 5,588,038,800 | 60% | 3,338,155,817 | 27.1 |
| GW | q | 8 | 39,872,032 | 5,980,804,800 | 59% | 3,528,765,680 | 28.7 |
| GW | q | 9 | 43,063,924 | 6,459,588,600 | 60% | 3,896,184,129 | 31.7 |
| GW | q | 10 | 59,353,224 | 8,902,983,600 | 56% | 5,005,146,389 | 40.7 |
| GW | q | 11 | 48,798,718 | 7,319,807,700 | 58% | 4,230,709,614 | 34.4 |
| GW | q | 12 | 51,587,534 | 7,738,130,100 | 59% | 4,530,553,062 | 36.8 |
| GW | q | 13 | 51,516,938 | 7,727,540,700 | 57% | 4,436,314,107 | 36.1 |
| GW | q | 14 | 52,368,526 | 7,855,278,900 | 57% | 4,491,431,340 | 36.5 |
| GW | q | 15 | 56,966,358 | 8,544,953,700 | 58% | 4,954,467,975 | 40.3 |
| GW | q | 16 | 55,542,004 | 8,331,300,600 | 58% | 4,855,542,511 | 39.5 |
| GW | q | 17 | 37,533,396 | 5,630,009,400 | 61% | 3,418,968,219 | 27.8 |
| GW | q | 18 | 36,991,832 | 5,548,774,800 | 62% | 3,456,767,923 | 28.1 |
| GW | q | 19 | 31,618,250 | 4,742,737,500 | 64% | 3,024,776,393 | 24.6 |
| GW | q | 20 | 38,664,584 | 5,799,687,600 | 61% | 3,552,969,635 | 28.9 |
| GW | q | 21 | 28,218,006 | 4,232,700,900 | 64% | 2,693,671,519 | 21.9 |
| GW | q | 22 | 26,671,494 | 4,000,724,100 | 66% | 2,621,832,865 | 21.3 |
| GW | q | 23 | 34,554,014 | 5,183,102,100 | 63% | 3,244,157,121 | 26.4 |
| GW | q | 24 | 30,864,458 | 4,629,668,700 | 64% | 2,963,367,536 | 24.1 |
| GW | q | 25 | 32,287,078 | 4,843,061,700 | 65% | 3,153,497,773 | 25.6 |
| GW | q | 26 | 32,492,524 | 4,873,878,600 | 64% | 3,142,976,513 | 25.6 |
| HM | A | 1 | 13,050,470 | 2,811,313,870 | 87% | 395,126,987 | 3.2 |
| HM | A | 2 | 12,050,092 | 2,486,855,227 | 87% | 365,133,969 | 3.0 |
| HM | A | 3 | 14,947,154 | 3,237,372,975 | 87% | 457,663,615 | 3.7 |
| HM | A | 4 | 11,685,630 | 2,540,352,532 | 89% | 362,727,598 | 2.9 |
| HM | A | 5 | 9,663,488 | 2,213,341,626 | 92% | 309,897,871 | 2.5 |
| HM | A | 6 | 11,026,316 | 2,475,282,758 | 89% | 344,552,640 | 2.8 |
| HM | A | 7 | 10,930,722 | 2,426,832,911 | 90% | 344,022,180 | 2.8 |
| HM | A | 8 | 11,923,520 | 2,729,913,087 | 88% | 368,147,079 | 3.0 |
| HM | A | 9 | 8,810,198 | 2,014,203,194 | 89% | 275,430,881 | 2.2 |
| HM | B | 1 | 6,873,724 | 1,592,507,642 | 89% | 212,937,256 | 1.7 |
| HM | B | 2 | 10,478,754 | 2,321,138,262 | 86% | 314,001,653 | 2.6 |
| HM | B | 3 | 9,072,450 | 2,102,879,818 | 90% | 284,420,111 | 2.3 |
| HM | B | 4 | 9,358,218 | 2,113,342,678 | 90% | 295,489,180 | 2.4 |
| HM | B | 5 | 14,281,626 | 3,226,827,968 | 87% | 433,950,892 | 3.5 |
| HM | B | 6 | 12,217,560 | 2,562,882,558 | 85% | 365,192,158 | 3.0 |
| HM | B | 7 | 11,966,220 | 2,749,143,195 | 88% | 366,838,009 | 3.0 |
| HM | C | 1 | 14,580,792 | 3,062,321,031 | 85% | 431,787,071 | 3.5 |
| HM | C | 2 | 15,787,984 | 3,517,350,681 | 85% | 472,100,197 | 3.8 |
| HM | C | 3 | 8,934,870 | 1,832,486,366 | 86% | 268,294,579 | 2.2 |
| HM | C | 4 | 8,201,794 | 1,846,396,810 | 89% | 255,294,330 | 2.1 |
| HM | C | 5 | 13,025,270 | 2,819,374,263 | 88% | 401,666,109 | 3.3 |
| HM | C | 6 | 17,085,714 | 3,734,936,693 | 84% | 500,886,416 | 4.1 |
| HM | C | 7 | 12,696,716 | 2,886,694,469 | 87% | 384,423,333 | 3.1 |
| HM | C | 8 | 11,099,246 | 2,601,743,388 | 86% | 334,313,700 | 2.7 |
| HM | C | 9 | 11,798,778 | 2,629,174,592 | 86% | 356,747,315 | 2.9 |
| HM | C | 10 | 12,960,568 | 2,838,217,781 | 85% | 385,073,962 | 3.1 |
| HM | C | 11 | 9,567,050 | 2,151,290,204 | 87% | 291,904,128 | 2.4 |
| HM | C | 12 | 7,205,790 | 1,632,042,851 | 86% | 217,136,672 | 1.8 |
| HM | C | 13 | 8,582,326 | 1,891,655,771 | 89% | 266,064,993 | 2.2 |
| HM | C | 14 | 9,292,548 | 1,991,951,781 | 88% | 287,735,401 | 2.3 |
| HM | C | 15 | 8,306,372 | 1,751,108,671 | 86% | 251,269,350 | 2.0 |
| HM | C | 16 | 12,657,764 | 2,684,632,268 | 87% | 384,881,062 | 3.1 |
| HM | C | 17 | 7,144,948 | 1,533,333,854 | 89% | 223,659,517 | 1.8 |
| HM | C | 18 | 11,568,066 | 2,493,366,900 | 85% | 344,565,512 | 2.8 |
| HM | C | 19 | 11,023,660 | 2,388,242,222 | 87% | 334,065,393 | 2.7 |
| HM | C | 20 | 9,750,124 | 2,171,361,549 | 87% | 297,763,928 | 2.4 |
| HM | D | 1 | 66,119,484 | 9,917,922,600 | 54% | 5,393,998,102 | 43.9 |
| HM | D | 2 | 57,666,440 | 8,649,966,000 | 47% | 4,096,911,959 | 33.3 |
| HM | D | 3 | 62,490,678 | 9,373,601,700 | 55% | 5,154,064,847 | 41.9 |
| HM | D | 4 | 52,453,766 | 7,868,064,900 | 59% | 4,613,577,509 | 37.5 |
| HM | D | 5 | 60,184,306 | 9,027,645,900 | 55% | 4,990,163,838 | 40.6 |
| HM | D | 6 | 52,625,776 | 7,893,866,400 | 57% | 4,511,503,360 | 36.7 |
| HM | D | 7 | 63,674,292 | 9,551,143,800 | 55% | 5,219,623,501 | 42.4 |
| HM | D | 8 | 56,169,480 | 8,425,422,000 | 56% | 4,757,465,892 | 38.7 |
| HM | E | 1 | 42,910,348 | 6,436,552,200 | 62% | 3,984,356,166 | 32.4 |
| HM | E | 2 | 40,184,186 | 6,027,627,900 | 63% | 3,802,671,756 | 30.9 |
| HM | E | 3 | 44,027,242 | 6,604,086,300 | 50% | 3,285,903,708 | 26.7 |
| HM | E | 4 | 38,295,782 | 5,744,367,300 | 56% | 3,237,762,049 | 26.3 |
| HM | E | 5 | 38,993,038 | 5,848,955,700 | 58% | 3,401,220,058 | 27.7 |
| HM | E | 6 | 39,685,254 | 5,952,788,100 | 62% | 3,715,020,521 | 30.2 |
| HM | E | 7 | 36,264,848 | 5,439,727,200 | 37% | 2,034,292,445 | 16.5 |
| HM | E | 8 | 33,857,042 | 5,078,556,300 | 44% | 2,240,546,181 | 18.2 |
| HM | E | 9 | 31,980,726 | 4,797,108,900 | 62% | 2,978,197,684 | 24.2 |
| HM | E | 10 | 29,756,000 | 4,463,400,000 | 63% | 2,813,977,919 | 22.9 |
| HM | F | 1 | 27,434,624 | 4,115,193,600 | 65% | 2,680,387,754 | 21.8 |
| HM | F | 2 | 31,506,698 | 4,726,004,700 | 67% | 3,165,219,164 | 25.7 |
| HM | G | 1 | 23,234,878 | 6,970,000,000 | 61% | 4,274,123,726 | 34.7 |
| HM | G | 2 | 21,386,665 | 6,416,000,000 | 64% | 4,094,504,032 | 33.3 |
| HM | G | 3 | 44,812,627 | 13,444,000,000 | 54% | 7,226,202,283 | 58.7 |
| HM | G | 4 | 21,947,537 | 6,584,000,000 | 62% | 4,084,963,082 | 33.2 |
| HM | G | 5 | 20,775,243 | 6,233,000,000 | 61% | 3,816,847,941 | 31.0 |
| HM | G | 6 | 20,412,166 | 6,124,000,000 | 63% | 3,864,869,138 | 31.4 |
| HM | G | 7 | 20,650,106 | 6,195,000,000 | 63% | 3,924,387,505 | 31.9 |
| HM | G | 8 | 22,438,067 | 6,731,000,000 | 63% | 4,245,594,586 | 34.5 |
| HM | G | 9 | 23,931,478 | 7,179,000,000 | 62% | 4,428,756,563 | 36.0 |
| HM | G | 10 | 23,596,951 | 7,079,000,000 | 58% | 4,083,773,009 | 33.2 |
| HM | G | 11 | 18,841,053 | 5,652,000,000 | 42% | 2,360,895,905 | 19.2 |
| HM | G | 12 | 22,694,920 | 6,808,000,000 | 63% | 4,296,419,755 | 34.9 |
| HM | G | 13 | 23,205,889 | 6,962,000,000 | 61% | 4,217,489,788 | 34.3 |
| HM | G | 14 | 16,000,308 | 4,800,000,000 | 50% | 2,385,186,208 | 19.4 |
| HM | G | 15 | 13,051,301 | 3,915,000,000 | 51% | 1,993,490,662 | 16.2 |
| HM | G | 16 | 19,029,795 | 5,709,000,000 | 65% | 3,703,282,614 | 30.1 |
| HM | G | 17 | 22,209,063 | 6,663,000,000 | 62% | 4,142,396,877 | 33.7 |
| HM | G | 18 | 21,593,699 | 6,478,000,000 | 64% | 4,118,524,707 | 33.5 |
| HM | G | 19 | 20,216,366 | 6,065,000,000 | 64% | 3,909,284,753 | 31.8 |
| HM | H | 1 | 79,428,636 | 11,993,724,036 | 78% | 9,340,612,034 | 75.9 |
| HM | H | 2 | 81,876,420 | 12,363,339,420 | 76% | 9,350,562,885 | 76.0 |
| HM | H | 3 | 83,257,466 | 12,571,877,366 | 79% | 9,887,821,549 | 80.4 |
| HM | H | 4 | 96,052,096 | 14,503,866,496 | 76% | 10,992,964,021 | 89.4 |
| HM | H | 5 | 163,813,294 | 24,735,807,394 | 70% | 17,324,774,384 | 140.9 |
| HM | H | 6 | 75,616,098 | 11,418,030,798 | 76% | 8,698,064,995 | 70.7 |
| HM | H | 7 | 73,480,260 | 11,095,519,260 | 75% | 8,366,580,113 | 68.0 |
| HM | H | 8 | 123,874,628 | 18,705,068,828 | 71% | 13,315,786,663 | 108.3 |
| HM | H | 9 | 80,328,866 | 12,129,658,766 | 74% | 9,021,456,376 | 73.3 |
| HM | H | 10 | 92,463,662 | 13,962,012,962 | 76% | 10,630,084,301 | 86.4 |
| HM | H | 11 | 84,850,168 | 12,812,375,368 | 74% | 9,427,831,376 | 76.6 |
| HM | H | 12 | 76,307,390 | 11,522,415,890 | 74% | 8,508,829,983 | 69.2 |
| HM | H | 13 | 71,073,642 | 10,732,119,942 | 79% | 8,437,785,462 | 68.6 |
| HM | H | 14 | 71,160,176 | 10,745,186,576 | 79% | 8,528,722,186 | 69.3 |
| HM | H | 15 | 67,944,082 | 10,259,556,382 | 81% | 8,293,521,401 | 67.4 |
| HM | H | 16 | 101,970,706 | 15,397,576,606 | 75% | 11,538,139,738 | 93.8 |
| HM | H | 17 | 97,648,608 | 14,744,939,808 | 75% | 11,111,701,134 | 90.3 |
| HM | H | 18 | 75,104,934 | 11,340,845,034 | 78% | 8,791,324,209 | 71.5 |
| HM | H | 19 | 75,022,554 | 11,328,405,654 | 76% | 8,648,985,863 | 70.3 |
| HM | H | 20 | 58,166,574 | 8,783,152,674 | 82% | 7,164,079,405 | 58.2 |
| HM | I | 1 | 72,187,770 | 10,828,165,500 | 51% | 5,471,176,168 | 44.5 |
| HM | I | 2 | 67,469,204 | 10,120,380,600 | 50% | 5,104,047,775 | 41.5 |
| HM | I | 3 | 65,568,588 | 9,835,288,200 | 48% | 4,730,902,085 | 38.5 |
| HM | I | 4 | 71,478,798 | 10,721,819,700 | 47% | 5,063,695,961 | 41.2 |
| HM | I | 5 | 143,958,708 | 21,593,806,200 | 45% | 9,627,237,676 | 78.3 |
| HM | I | 6 | 74,049,188 | 11,107,378,200 | 50% | 5,595,956,997 | 45.5 |
| HM | J | 1 | 119,954,864 | 18,113,184,464 | 27% | 4,827,092,610 | 39.2 |
| HM | J | 2 | 57,759,354 | 8,721,662,454 | 53% | 4,596,599,541 | 37.4 |
| HM | J | 3 | 56,835,130 | 8,582,104,630 | 56% | 4,784,946,996 | 38.9 |
| HM | J | 4 | 48,704,852 | 7,354,432,652 | 56% | 4,130,376,063 | 33.6 |
| HM | J | 5 | 54,355,392 | 8,207,664,192 | 74% | 6,096,149,221 | 49.6 |
| HM | K | 1 | 53,908,848 | 8,140,236,048 | 56% | 4,575,453,016 | 37.2 |
| HM | K | 2 | 54,821,824 | 8,278,095,424 | 40% | 3,330,449,761 | 27.1 |
| HM | K | 3 | 56,730,052 | 8,566,237,852 | 33% | 2,790,265,236 | 22.7 |
| HM | K | 4 | 55,425,242 | 8,369,211,542 | 45% | 3,800,136,585 | 30.9 |
| HM | K | 5 | 50,924,844 | 7,689,651,444 | 77% | 5,924,618,179 | 48.2 |
| HM | L | 1 | 49,690,112 | 7,503,206,912 | 77% | 5,774,309,111 | 46.9 |
| HM | L | 2 | 55,670,620 | 8,406,263,620 | 78% | 6,532,991,698 | 53.1 |
| HM | L | 3 | 54,636,116 | 8,250,053,516 | 76% | 6,238,959,949 | 50.7 |
| HM | L | 4 | 57,314,972 | 8,654,560,772 | 54% | 4,710,819,414 | 38.3 |
| HM | L | 5 | 52,713,520 | 7,959,741,520 | 57% | 4,529,326,265 | 36.8 |
| HM | M | 1 | 45,752,014 | 6,908,554,114 | 60% | 4,162,691,653 | 33.8 |
| HM | M | 2 | 47,904,106 | 7,233,520,006 | 83% | 5,984,401,985 | 48.7 |
| HM | M | 3 | 57,213,898 | 8,639,298,598 | 54% | 4,635,688,984 | 37.7 |
| HM | M | 4 | 55,332,950 | 8,355,275,450 | 57% | 4,802,184,298 | 39.0 |
| HM | M | 5 | 54,391,828 | 8,213,166,028 | 75% | 6,121,467,501 | 49.8 |
| HM | N | 1 | 61,583,298 | 9,299,077,998 | 27% | 2,550,697,797 | 20.7 |
| HM | N | 2 | 50,921,896 | 7,689,206,296 | 81% | 6,243,929,352 | 50.8 |
| HM | N | 3 | 53,686,292 | 8,106,630,092 | 81% | 6,576,902,571 | 53.5 |
| HM | N | 4 | 57,091,806 | 8,620,862,706 | 57% | 4,932,765,991 | 40.1 |
| HM | N | 5 | 568,797,242 | 85,888,383,542 | 20% | 17,080,837,742 | 138.9 |
| HM | O | 1 | 52,478,650 | 7,924,276,150 | 55% | 4,394,488,368 | 35.7 |
| HM | O | 2 | 769,547,018 | 116,201,599,718 | 22% | 25,504,682,161 | 207.4 |
| HM | O | 3 | 53,010,852 | 8,004,638,652 | 58% | 4,624,285,714 | 37.6 |
| HM | O | 4 | 57,318,330 | 8,655,067,830 | 31% | 2,718,322,774 | 22.1 |
| HM | O | 5 | 56,675,410 | 8,557,986,910 | 54% | 4,592,131,384 | 37.3 |
| HM | P | 1 | 53,457,372 | 8,072,063,172 | 58% | 4,709,123,438 | 38.3 |
| HM | P | 2 | 264,626,282 | 39,958,568,582 | 26% | 10,254,233,045 | 83.4 |
| HM | P | 3 | 52,669,190 | 7,953,047,690 | 58% | 4,607,430,284 | 37.5 |
| HM | P | 4 | 54,321,420 | 8,202,534,420 | 55% | 4,551,627,892 | 37.0 |
| HM | P | 5 | 58,237,198 | 8,793,816,898 | 53% | 4,666,835,704 | 37.9 |
| HM | Q | 1 | 47,050,980 | 7,104,697,980 | 81% | 5,771,512,861 | 46.9 |
| HM | Q | 2 | 57,268,148 | 8,647,490,348 | 55% | 4,783,824,290 | 38.9 |
| HM | Q | 3 | 54,039,968 | 8,160,035,168 | 53% | 4,335,249,147 | 35.2 |
| HM | Q | 4 | 56,475,846 | 8,527,852,746 | 57% | 4,840,088,279 | 39.4 |
| HM | Q | 5 | 657,548,228 | 99,289,782,428 | 22% | 21,790,998,458 | 177.2 |
| HM | R | 1 | 55,173,006 | 8,331,123,906 | 57% | 4,749,643,904 | 38.6 |
| HM | R | 2 | 47,809,986 | 7,219,307,886 | 69% | 4,953,143,171 | 40.3 |
| HM | R | 3 | 57,263,084 | 8,646,725,684 | 56% | 4,809,966,042 | 39.1 |
| HM | R | 4 | 56,877,358 | 8,588,481,058 | 54% | 4,672,709,361 | 38.0 |
| HM | R | 5 | 834,901,790 | 126,070,170,290 | 21% | 26,539,072,886 | 215.8 |
| YM | U | 1 | 17,554,310 | 3,916,010,892 | 87% | 532,137,484 | 4.3 |
| YM | U | 2 | 17,347,782 | 3,933,418,815 | 87% | 528,971,686 | 4.3 |
| YM | U | 3 | 13,753,778 | 3,064,230,933 | 87% | 420,833,194 | 3.4 |
| YM | U | 4 | 17,663,514 | 3,958,223,025 | 88% | 544,310,783 | 4.4 |
| YM | U | 5 | 14,445,180 | 3,203,584,978 | 85% | 428,825,338 | 3.5 |
| YM | U | 6 | 15,982,784 | 3,589,973,983 | 86% | 483,877,126 | 3.9 |
| YM | U | 7 | 17,635,184 | 3,964,396,649 | 82% | 506,587,543 | 4.1 |
| YM | V | 1 | 10,558,244 | 2,115,058,767 | 86% | 319,141,684 | 2.6 |
| YM | V | 2 | 11,246,744 | 2,375,906,382 | 87% | 344,282,327 | 2.8 |
| YM | V | 3 | 11,020,452 | 2,306,616,994 | 90% | 346,581,873 | 2.8 |
| YM | V | 4 | 10,780,966 | 2,296,630,203 | 86% | 325,841,715 | 2.6 |
| YM | V | 5 | 13,544,796 | 2,772,485,059 | 88% | 415,695,937 | 3.4 |
| YM | V | 6 | 12,947,332 | 2,959,174,103 | 86% | 387,540,348 | 3.2 |
| YM | V | 7 | 15,368,476 | 2,982,045,259 | 89% | 477,914,507 | 3.9 |
| YM | V | 8 | 11,366,038 | 2,534,337,706 | 87% | 347,374,720 | 2.8 |
| YM | V | 9 | 9,936,942 | 2,199,329,310 | 89% | 308,867,310 | 2.5 |
| YM | V | 10 | 8,567,482 | 2,029,766,667 | 90% | 269,404,058 | 2.2 |
| CH | Z | 1 | 79,762,778 | 12,044,179,478 | 79% | 4,831,711,574 | 39.3 |
| CH | Z | 2 | 68,669,652 | 10,369,117,452 | 81% | 4,222,026,165 | 34.3 |
| CH | Z | 3 | 76,432,970 | 11,541,378,470 | 78% | 4,661,574,270 | 37.9 |
| CH | Z | 4 | 81,960,384 | 12,376,017,984 | 78% | 9,602,493,541 | 78.1 |

\*1 GW, animals derived from Onagawa; HM, animals derived from Onahama; YM, animals derived from Okayama; KS, animals derived from Kisarazu
\*2 These percentages were estimated using the FASTQC program
\*3 Depth was calculated under the assumption that the genome size is 123 Mb (Satou et al., 2019 Genome Biol. Evol. 11:3144–3157 doi:10.1093/gbe/evz228).

**Table S2.** Heterozygosity rates of animals in the closed colonies.

| Origin | Sample Group | IDs | Heterozygosity rate |
| --- | --- | --- | --- |
| GW | e | 1 | 0.85% |
| GW | e | 2 | 0.85% |
| GW | e | 3 | 0.30% |
| GW | e | 4 | 0.86% |
| GW | e | 5 | 0.84% |
| GW | e | 6 | 0.83% |
| GW | e | 7 | 0.84% |
| GW | e | 8 | 0.85% |
| GW | e | 9 | 0.84% |
| GW | e | 10 | 0.86% |
| GW | g | 1 | 0.88% |
| GW | g | 2 | 0.89% |
| GW | g | 3 | 0.85% |
| GW | g | 4 | 0.89% |
| GW | g | 5 | 0.87% |
| GW | g | 6 | 0.90% |
| GW | g | 7 | 0.90% |
| GW | g | 8 | 0.90% |
| GW | g | 9 | 0.89% |
| GW | g | 10 | 0.87% |
| GW | g | 11 | 0.87% |
| GW | g | 12 | 0.87% |
| GW | g | 13 | 0.85% |
| GW | g | 14 | 0.89% |
| GW | p | 1 | 0.82% |
| GW | p | 2 | 0.90% |
| GW | p | 3 | 0.92% |
| GW | p | 4 | 0.91% |
| GW | p | 5 | 0.87% |
| GW | p | 6 | 0.91% |
| GW | p | 7 | 0.89% |
| GW | p | 8 | 0.81% |
| GW | p | 9 | 0.90% |
| GW | p | 10 | 0.91% |
| GW | p | 11 | 0.86% |
| GW | p | 12 | 0.93% |
| GW | p | 13 | 0.81% |
| GW | p | 14 | 0.91% |
| GW | p | 15 | 0.91% |
| GW | p | 16 | 0.90% |
| GW | p | 17 | 0.88% |
| GW | p | 18 | 0.86% |
| GW | p | 19 | 0.89% |
| GW | p | 20 | 0.92% |
| GW | p | 21 | 0.82% |
| GW | p | 22 | 0.90% |
| GW | q | 1 | 0.82% |
| GW | q | 2 | 0.85% |
| GW | q | 3 | 0.80% |
| GW | q | 4 | 0.75% |
| GW | q | 5 | 0.82% |
| GW | q | 6 | 0.79% |
| GW | q | 7 | 0.78% |
| GW | q | 8 | 0.83% |
| GW | q | 9 | 0.80% |
| GW | q | 10 | 0.85% |
| GW | q | 11 | 0.83% |
| GW | q | 12 | 0.79% |
| GW | q | 13 | 0.81% |
| GW | q | 14 | 0.86% |
| GW | q | 15 | 0.79% |
| GW | q | 16 | 0.79% |
| GW | q | 17 | 0.77% |
| GW | q | 18 | 0.79% |
| GW | q | 19 | 0.77% |
| GW | q | 20 | 0.83% |
| GW | q | 21 | 0.80% |
| GW | q | 22 | 0.73% |
| GW | q | 23 | 0.83% |
| GW | q | 24 | 0.82% |
| GW | q | 25 | 0.80% |
| GW | q | 26 | 0.85% |
| HM | E | 1 | 0.87% |
| HM | E | 2 | 0.86% |
| HM | E | 3 | 0.84% |
| HM | E | 4 | 0.83% |
| HM | E | 5 | 0.80% |
| HM | E | 6 | 0.86% |
| HM | E | 7 | 0.44% |
| HM | E | 8 | 0.53% |
| HM | E | 9 | 0.82% |
| HM | E | 10 | 0.81% |
| HM | G | 1 | 0.95% |
| HM | G | 2 | 0.96% |
| HM | G | 3 | 0.97% |
| HM | G | 4 | 0.95% |
| HM | G | 5 | 0.94% |
| HM | G | 6 | 0.95% |
| HM | G | 7 | 0.94% |
| HM | G | 8 | 0.96% |
| HM | G | 9 | 0.97% |
| HM | G | 10 | 0.96% |
| HM | G | 11 | 0.60% |
| HM | G | 12 | 0.96% |
| HM | G | 13 | 0.95% |
| HM | G | 14 | 0.63% |
| HM | G | 15 | 0.70% |
| HM | G | 16 | 0.94% |
| HM | G | 17 | 0.95% |
| HM | G | 18 | 0.95% |
| HM | G | 19 | 0.94% |
| HM | H | 1 | 0.85% |
| HM | H | 2 | 0.85% |
| HM | H | 3 | 0.83% |
| HM | H | 4 | 0.87% |
| HM | H | 5 | 0.85% |
| HM | H | 6 | 0.86% |
| HM | H | 7 | 0.86% |
| HM | H | 8 | 0.92% |
| HM | H | 9 | 0.92% |
| HM | H | 10 | 0.91% |
| HM | H | 11 | 0.88% |
| HM | H | 12 | 0.83% |
| HM | H | 13 | 0.86% |
| HM | H | 14 | 0.88% |
| HM | H | 15 | 0.82% |
| HM | H | 16 | 0.93% |
| HM | H | 17 | 0.88% |
| HM | H | 18 | 0.89% |
| HM | H | 19 | 0.91% |
| HM | H | 20 | 0.87% |
| HM | I | 1 | 1.19% |
| HM | I | 2 | 1.19% |
| HM | I | 3 | 0.83% |
| HM | I | 4 | 0.64% |
| HM | I | 5 | 0.75% |
| HM | I | 6 | 0.87% |

## Notes

### Competing Interest Statement

The authors have declared no competing interest.

